# Ankyrin-G Regulates Forebrain Connectivity and Network Synchronization via Interaction with GABARAP

**DOI:** 10.1101/307512

**Authors:** AD Nelson, RN Caballero-Florán, JC Rodríguez Díaz, J Li, K Chen, KK Walder, V Bennett, LF Lopez-Santiago, MG McInnis, LL Isom, C Wang, M Zhang, KS Jones, PM Jenkins

## Abstract

GABAergic circuits are critical for the synchronization and higher order function of brain networks, and defects in this circuitry are linked to neuropsychiatric diseases, including bipolar disorder, schizophrenia, and autism. Work in cultured neurons has shown that ankyrin-G plays a key role in the regulation of GABAergic synapses on the axon initial segment and somatodendritic domain of pyramidal neurons where it interacts directly with the GABAA receptor associated protein (GABARAP) to stabilize cell surface GABAA receptors. Here, we generated a knock-in mouse model expressing a mutation that abolishes the ankyrin-G/GABARAP interaction (*Ank3* W1989R) to understand how ankyrin-G and GABARAP regulate GABAergic circuitry *in vivo*. We found that *Ank3* W1989R mice exhibit a striking reduction in forebrain GABAergic synapses resulting in pyramidal cell hyperexcitability and disruptions in network synchronization. In addition, we identified changes in pyramidal cell dendritic spines and axon initial segments consistent with compensation for hyperexcitability. Finally, we identified the *ANK3* W1989R variant in a family with bipolar disorder, suggesting a potential role of this variant in disease. Our results highlight the importance of ankyrin-G in regulating forebrain circuitry and provide novel insights into how *ANK3* loss-of-function variants may contribute to human disease.

## INTRODUCTION

GABAergic interneurons are essential for the proper synchronization and function of neuronal networks that underlie normal cognition, mood, and behavior. GABAergic interneurons target to unique postsynaptic domains on excitatory neurons; however, the molecular mechanisms underlying the subcellular organization of cortical GABAergic synapses remain poorly understood. Abnormalities in GABAergic interneuron circuitry and decreased gamma oscillations have been implicated in many neurodevelopmental and neuropsychiatric disorders^1–8^. Thus, the understanding of the cellular and molecular mechanisms that contribute to the development and function of GABAergic synapses as well as identification of genetic variants that contribute to neuropsychiatric disorders is critical to the discovery of new therapeutic agents for the treatment of diseases involving altered inhibitory circuits.

*ANK3* encodes ankyrin-G, a fundamental scaffolding protein that organizes critical plasma membrane domains^9, 10^. Alternative splicing of *ANK3* in the brain gives rise to three main isoforms of ankyrin-G: the canonical 190 kDa isoform, a 270 kDa isoform, and a giant, 480 kDa isoform. The 190 kDa isoform is expressed in most tissues and cell types throughout the body including brain, heart, skeletal muscle, kidney, and retina. The 270 kDa and 480 kDa isoforms of ankyrin-G are predominantly expressed in the nervous system, and arise from alternative splicing of a 7.8 kb vertebrate-specific exon^9, 11^. The 480 kDa ankyrin-G isoform has been identified as the master organizer of axon initial segments (AIS) and nodes of Ranvier, sites of action potential (AP) initiation and propagation^10^. This splice variant is necessary for the proper clustering of voltage-gated sodium channels, KCNQ2/3 potassium channels, the cell adhesion molecule neurofascin-186, and the cytoskeletal protein βIV-spectrin to excitable domains (reviewed in ^12^).

Importantly, the 480 kDa ankyrin-G isoform has also been shown to stabilize GABAergic synapses on the soma and AIS of excitatory pyramidal neurons by interacting with the GABAA receptor-associated protein (GABARAP) to inhibit GABAA-receptor endocytosis ^13^. GABARAP and GABARAP-like 1, members of the ubiquitin-like LC3 family of microtubule-associated proteins, mediate GABAA-receptor trafficking between the cell surface and intracellular compartments^14^. GABARAP and other members of the LC3 family interact with LC3-interacting region (LIR) motifs^15^. The giant exon that encodes the 480 kDa ankyrin-G isoform includes an LIR motif, which includes residue W1989^13, 16^. Mutation of W1989 to arginine (W1989R) completely abolished the binding between ankyrin-G and GABARAP^13^. Deletion of wild-type (WT) ankyrin-G and replacement with W1989R 480 kDa ankyrin-G failed to rescue GABAA-receptors to the soma and AIS or restore miniature inhibitory post-synaptic currents (mIPSCs) in cultured mouse hippocampal neurons^13^. Taken together, these findings suggested that 480 kDa ankyrin-G plays a critical role in stabilizing GABAergic synapses *in vitro*; however, whether ankyrin-G loss-of-function at GABAergic synapses disrupts forebrain circuitry *in vivo* has not been investigated.

Here, we have generated a novel knock-in mouse model expressing *Ank3* W1989R. This allowed us to study, for the first time, the relationship between the 480 kDa ankyrin-G isoform and GABAergic synapse formation and function *in vivo* in a model that survives to adulthood and is capable of forming the AIS and nodes of Ranvier. We show that the *Ank3* W1989R mutation causes decreases in GABAergic synapses in layer II/III of somatosensory cortex and CA1 of hippocampus, while sparing inhibitory synapses on cerebellar Purkinje neurons. The decreases in inhibitory synapses cause hyperexcitability of cortical and hippocampal pyramidal neurons and decreases in gamma oscillations. Interestingly, we also detect changes consistent with compensation for the loss of inhibitory tone, including shortening of the AIS and decreases in dendritic spine density and excitatory postsynaptic currents. Finally, we report the identification of a family with bipolar disorder (BD) that carries the *Ank3* W1989R human variant (10:61834674 A/G), which may contribute to the pathophysiology of psychiatric disease.

## RESULTS

### *Ank3* W1989, located within the giant exon of ankyrin-G, is necessary for binding to a hydrophobic pocket in GABARAP

The 480 kDa *Ank3* splice variant interacts with GABARAP to inhibit GABAA-receptor endocytosis and stabilize GABAergic synapses^13^. Here, we explored the molecular basis governing this interaction by resolving the crystal structure of the ankyrin-G/GABARAP complex. Crystallography data show that the LIR motif within the giant exon of ankyrin-G contains aromatic residues, W1989 and F1992, which insert into two hydrophobic pockets of GABARAP (Fig. 1A and Supplementary Fig. 1A). Moreover, a unique C-terminal helix extension contributes to ankyrin-G/GABARAP binding by forming a critical salt bridge between residues E1996 of ankyrin-G and R67 of GABARAP and additional hydrophobic interactions with a hydrophobic surface of GABARAP (Supplementary Fig. 1A). This newly defined binding mode with the presence of the C-terminal helix is unique compared to previously known GABARAP/LIR motifs or other LC3 family members/LIR interactions (Fig. 1B), suggesting a specific neuronal function of the ankyrin-G/GABARAP interaction outside the autophagic processes for GABARAP^17^. To address the role of the W1989 residue in more detail, we performed Isothermal Titration Calorimetry (ITC) to quantitatively measure the dissociation constant (K_d_) between a series of ankyrin-G truncations and GABARAP. Using this approach, we mapped the minimal region of ankyrin-G that is capable of binding to GABARAP to a fragment of 26 amino acids (residues 1985-2010), which included residue W1989 contained within the canonical LIR motif (Supplementary Fig. 1B). This ankyrin-G fragment associated with GABARAP with a K_d_ of 2.9 nM, which is more than 1000-fold stronger than previously reported interactions between GABARAP and other LIR motifs (Supplementary Fig. 1C and Fig. 1B)^17^. Mutation of the W1989 residue to R abolished the interaction between ankyrin-G and GABARAP, and decreased the binding affinity to approximately 11 μM, which is 4000-fold weaker than WT ankyrin-G (Supplementary Fig.1C)^13^. Overall, these results demonstrate that ankyrin-G residue W1989 is necessary for high affinity binding to GABARAP, while F1992 and the C-terminal helix extension play important roles in maintaining this interaction.

**Fig. 1:**
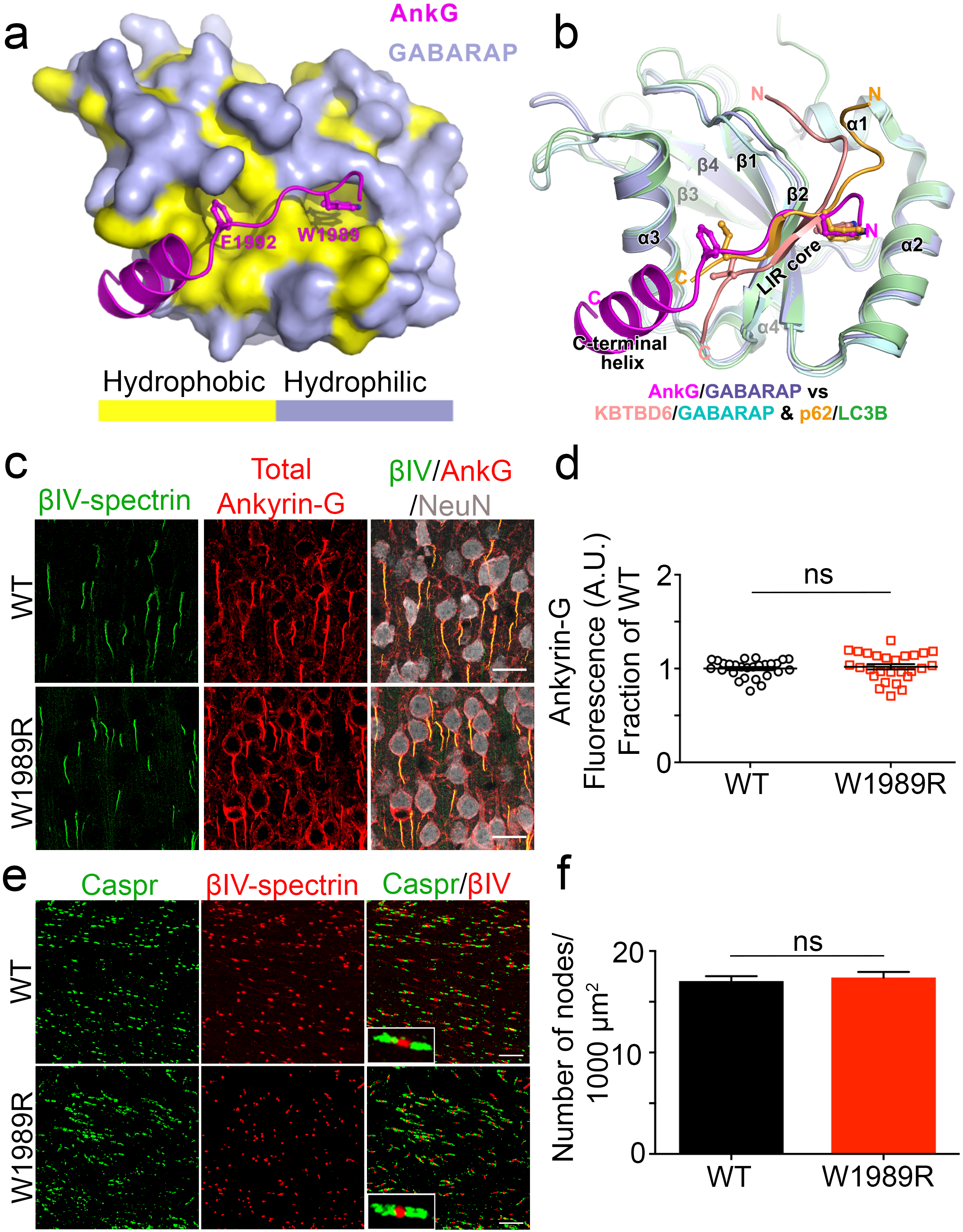
AIS and nodes of Ranvier are maintained in the *Ank3* W1989R mouse model. **(a)** Combined surface (GABARAP) and ribbon-stick model (ankyrin-G) showing a hydrophobic pocket of GABARAP is accommodated by the critical W1989 reside within the giant exon of ankyrin-G. The surface for hydrophobic residues of GABARAP are shown in yellow while the hydrophilic surfaces are light purple. The crystal structure of ankyrin-G/GABARAP complex is at a 2.2Å resolution. **(b)** Ribbon-stick model of superposition of ankyrin-G/GABARAP, KBTBD6/GABARAP and p62/LC3B complex structures showing the comparison of ankyrin-G/GABARAP complex with common binding mode of LIR/Atg8s. **(c)** Representative images from coronal sections of layer II/III somatosensory cortex of P30-35 WT (top) and *Ank3* W1989R homozygous (bottom) mice. Immunostaining for βIV-spectrin (green), total ankyrin-G (red), and NeuN (white). Scale bar: 20 μm. **(d)** Quantification of ankyrin-G fluorescence intensity (a.u.) as fraction of WT between WT (black circles) and *Ank3* W1989R homozygous (red squares) mice. *t-test* P = 0.5618 (WT: 1 ± 0.02, N=3, n=27; W1989R: 1.02 ± 0.03, N=3, n=27). **(e)** Representative images of nodes of Ranvier from corpus callosum of P30 WT (top) and *Ank3* W1989R homozygous (bottom) mice. Sections immunostained for the paranodal marker Caspr (green) and βIV-spectrin (red). Scale bar: 20μm. **(f)** Quantification of the total number of nodes of Ranvier per 1000 μm^2^ from WT (black bar) and *Ank3* W1989R homozygous (red bar) mice. *t-test* P = 0.62, ns: not significant (WT: 17.04 ± 0.5, N=3, n=20; W1989R: 17.4 ± 0.5, N=3, n=18). Data shown as mean ± SEM.

### W1989R 480 kDa ankyrin-G maintains WT functionality in assembly of the AIS and nodes of Ranvier *in vivo*

Several mouse models have been generated to evaluate the neuronal role of ankyrin-G *in vivo*, however these mice either die early in development or lack multiple cellular domains, making it difficult to understand the specific role of ankyrin-G-dependent GABAergic circuits^13, 16, 18^. To examine the role of 480 kDa ankyrin-G specifically in GABAergic synapse formation and function *in vivo*, we generated a knock-in mouse model expressing the W1989R (tgg>cgg) mutation within the giant exon of ankyrin-G. Homozygous *Ank3* W1989R mice survive well into adulthood, similar to WT littermate controls, living to at least P350. Homozygous *Ank3* W1989R mice are similar to WT mice in appearance and grooming behavior, and have no obvious neurological phenotype.

Previous studies demonstrated that acute deletion of WT ankyrin-G followed by rescue with W1989R 480 kDa ankyrin-G failed to restore GABAA-receptor clustering and mIPSCs in cultured hippocampal neurons. However, the W1989R mutant appropriately localized to the AIS and clustered all known binding partners^13^. Thus, we expected the knock-in *Ank3* W1989R mutation to function similar to WT at the AIS and nodes of Ranvier *in vivo*. As predicted, immunolabeling of cortical neurons in layer II/III of the somatosensory cortex in coronal brain sections from P30-35 mice with antibodies specific to ankyrin-G revealed that W1989R 480 kDa ankyrin-G appropriately localized to the AIS (Fig. 1C and D). In addition, W1989R 480 kDa ankyrin-G clustered all tested ankyrin-G binding partners to the AIS, including βIV spectrin, neurofascin, KCNQ2/3 channels, and voltage-gated sodium channels (Fig. 1C and Supplementary Fig. 2).

In addition to regulating the AIS, 480 kDa ankyrin-G plays a central role in the formation and maintenance of nodes of Ranvier^16^. Analysis of the corpus callosum in homozygous *Ank3* W1989R mice revealed no detectable changes in the total number of nodes of Ranvier (Fig. 1E and F) or nodal length (WT: 1.48 ± 0.04 μm, N=3, n=109; W1989R: 1.6 ± 0.03 μm, N=3, n=116) compared to WT mice (Fig. 1F). Thus, W1989R 480 kDa ankyrin-G maintains WT functionality in forming the AIS and nodes of Ranvier *in vivo*.

### GABAergic synapses and synaptic activity are decreased in *Ank3* W1989R forebrain pyramidal neurons *in vivo*

GABAergic interneurons synapse onto the dendrites, soma, and AIS of pyramidal neurons, and regulate excitability, synaptic transmission, and the synchronization of neuronal ensembles^19, 20^. To determine the effect of *Ank3* W1989R on GABAergic synapses *in vivo*, we immunostained WT and homozygous mutant P30-35 coronal brain sections with antibodies to the presynaptic inhibitory marker, vesicular GABA transporter (vGAT). *Ank3* W1989R mice showed a ~50-66% reduction in the number of GABAergic synapse clusters on the somatodendritic domain and AIS of cortical pyramidal neurons compared to WT neurons (Fig. 2A, B, and C). Moreover, clustering of postsynaptic GABAA-receptors was significantly decreased in dissociated hippocampal neurons from *Ank3* W1989R mice vs. WT (Supplementary Fig. 3A and B). These findings demonstrate that both pre- and post-synaptic structural components of GABAergic synapses are lost when 480 kDa ankyrin-G is unable to interact with GABARAP *in vivo*.

**Fig. 2:**
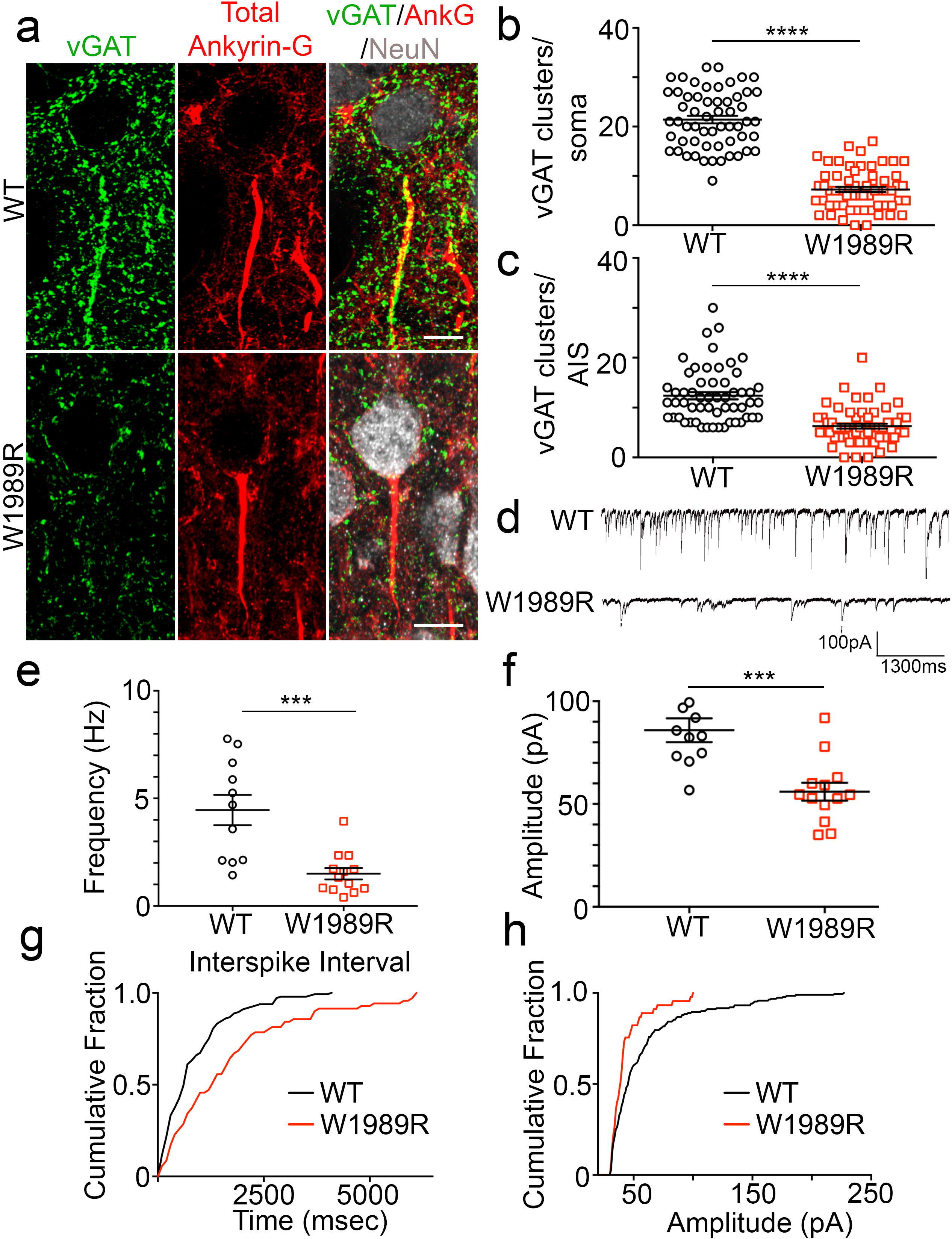
GABAergic synapses are reduced on the soma and AIS of *Ank3* W1989R pyramidal neurons. **(a)** Representative images of GABAergic synapses from layer II/III somatosensory cortex of P30 WT (top) and *Ank3* W1989R homozygous (bottom) mice. Coronal brain sections immunostained with a presynaptic GABAergic marker vGAT (green), total ankyrin-G (red), and NeuN (white). Scale bar: 20 μm. **(b)** Quantification of the total number of vGAT-positive clusters per soma above a set intensity threshold from WT (black circles) and *Ank3* W1989R homozygous (red squares) mice. *t-test* ****P < 0.0001 (WT: 21.42 ± 0.8, N=3, n=57; W1989R: 7.25 ± 0.5, N=3, n=60). **(c)** Quantification of total number of vGAT-positive clusters per AIS from WT (black circles) and *Ank3* W1989R homozygous (red squares) mice. *t-test* ****P < 0.0001 (WT: 12.38 ± 0.7, N=3, n=56; W1989R: 6.27 ± 0.5, N=3, n=60). **(d)** Spontaneous inhibitory post-synaptic current (sIPSC) representative traces from layer II/III somatosensory cortical neurons in WT and *Ank3* W1989R slices. Scale bars: 100 pA, 1300 ms. **(e)** Quantification of sIPSC frequency in WT (black circles) and *Ank3* W1989R (red squares) brain slices. *t-test* ***P = 0.0004 (WT: 4.5 ± 0.7 Hz, n=11; W1989R: 1.5 ± 0.3 Hz, n=13). **(f)** Quantification of sIPSC amplitude in WT (black circles) and *Ank3* W1989R (red squares) brain slices. *t-test* ***P = 0.0004 (WT: 85.9 ± 5.8 pA, n=11; W1989R: 56.0 ± 4.4 pA, n=13). **(g)** Representative cumulative histogram of interspike interval frequency of WT (black line) and *Ank3* W1989R homozygous (red line) sIPSCs in layer II/III somatosensory cortical neurons. *Kolmogorov-Smirnov* ****P < 0.0001. **(h)** Representative interspike interval amplitude of WT (black line) and *Ank3* W1989R homozygous (red line) sIPSCs. *Kolmogorov-Smirnov* ****P < 0.0001. Data shown as mean ± SEM.

To evaluate the functional consequences of *Ank3* W1989R on GABAergic signaling, we performed whole-cell patch-clamp recordings in acute brain slices of P25-48 WT and homozygous mutant mice. The frequency and amplitude of spontaneous inhibitory post-synaptic currents (sIPSCs) were significantly reduced in layer II/III cortical neurons as well as CA1 hippocampal neurons of *Ank3* W1989R mice relative to WT (Fig. 2D, E, F and Supplementary Fig. 3C, D, E). The magnitude of reduction in sIPSC frequency (~33%) and amplitude (~60%) were similar between pyramidal neurons in cortical layer II/III and CA1 hippocampal neurons in *Ank3* W1989R mice compared to WT, suggesting that the binding of 480 kDa ankyrin-G to GABARAP may be a common mechanism for stabilizing GABAergic synapses in the forebrain. Both layer II/III cortical neurons (Fig. 2D, G, and H) and CA1 hippocampal neurons (Supplementary Fig. 3C, F, and G) exhibited longer interspike intervals and the cumulative distribution of sIPSC amplitudes displayed smaller events in *Ank3* W1989R mice compared to WT. To address whether the decrease in quantal GABA release observed in *Ank3* W1989R mice is independent of AP firing, we measured mIPSCs in the presence of 1 μM tetrodotoxin (TTX) in acute brain slices of P25-48 mice. Consistent with the sIPSC data, we observed a significant reduction in both the frequency and amplitude of mIPSCs in *Ank3* W1989R cortical neurons versus WT (Supplementary Fig. 4A, B, C, and D). The iPSC response was completely attenuated following the administration of the GABAA-receptor antagonist bicuculline, confirming that the loss of inhibitory tone in *Ank3* W1989R neurons is specific to GABAA-receptor function rather than to other means of inhibitory synaptic transmission (Supplementary Fig. 4D).

**Fig. 3:**
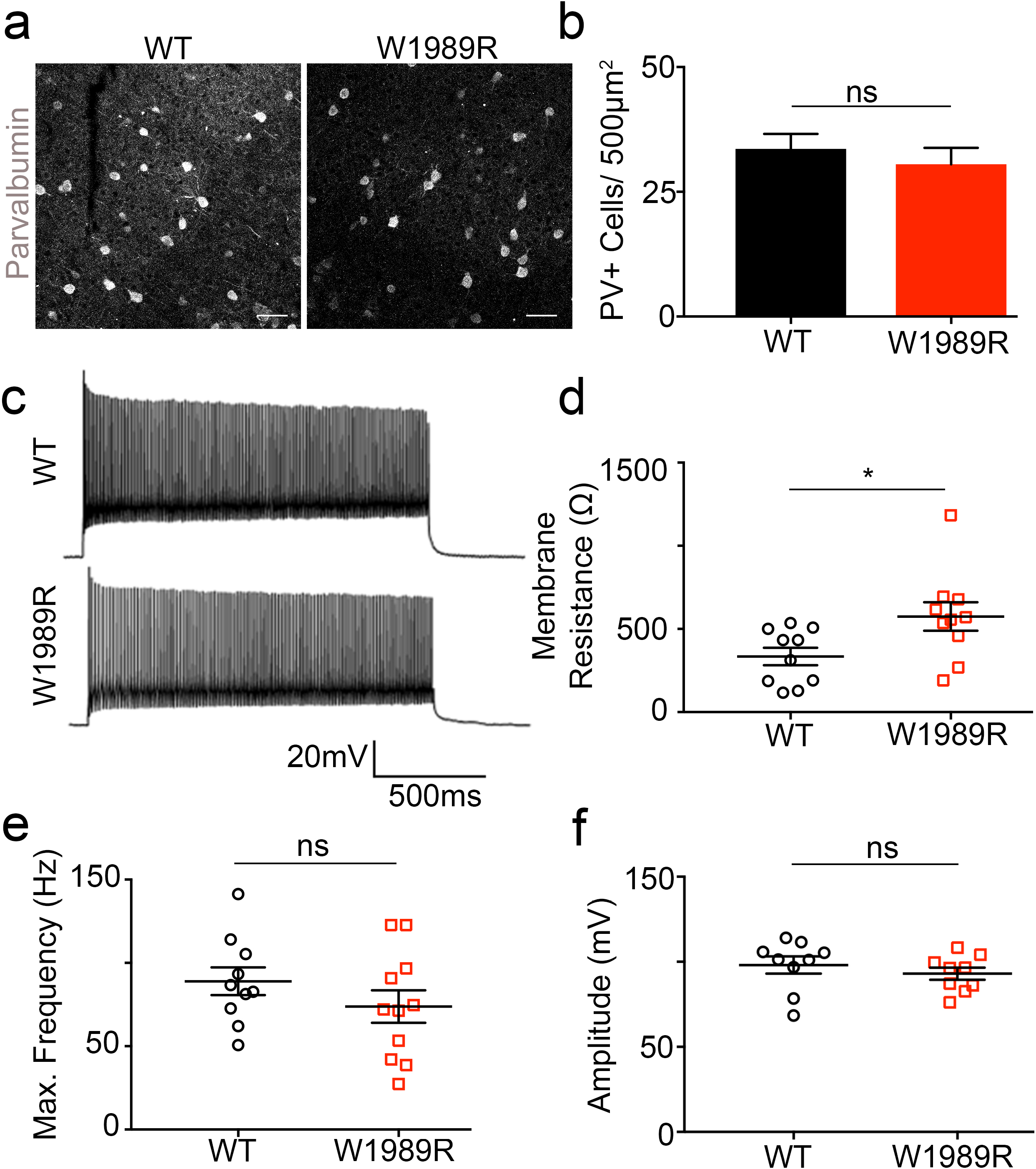
Normal density and fast-spiking properties of *Ank3* W1989R PV+ GABAergic interneurons. **(a)** Representative images of parvalbumin-positive (PV+) interneurons in layer II/III somatosensory cortex of P30 WT (left) and *Ank3* W1989R homozygous (right) mice. Coronal brain sections immunostained with PV (white). Scale bar: 50 μm. **(b)** Quantification of total number of PV-positive cells per 500 μm^2^ from WT (black bar) and *Ank3* W1989R homozygous (red bar) sections. *t-test* P = 0.497, ns, not significant (WT: 33.59 ± 3.0, N=3, n=15; W1989R: 30.53 ± 3.3, N=3, n=14). **(c)** Representative traces of evoked firing patterns and AP frequencies of fast-spiking PV+ interneurons in layer II/III somatosensory cortex from WT (top) and *Ank3* W1989R homozygous (bottom) brain slices. Scale bar: 500 ms. **(d)** Quantification of membrane resistance of PV+ cells in WT (black circles) and *Ank3* W1989R homozygous (red squares). *t-test* *P = 0.0276 (WT: 334.5 ± 52.8, n=10; W1989R: 574.9 ± 85.31, n=10). **(e)** Quantification of PV+ cell maximum frequency in WT (black circles) and *Ank3* W1989R homozygous (red squares). *t-test* P = 0.256 (WT: 88.99 ± 8.3, n=10; W1989R: 73.83 ± 9.7, n=11). **(f)** Quantification of single AP amplitude of PV+ in WT (black circles) and *Ank3* W1989R homozygous (red squares). *t-test* P = 0.422 (WT: 98.09 ± 5.0, n=9; W1989R: 93.02 ± 3.5, n=10). Data shown as mean ± SEM.

To determine whether the 480 kDa ankyrin-G interaction with GABARAP is a general mechanism for stabilizing GABAergic synapses, we immunostained Purkinje neurons in the cerebellum of sagittal brain sections from P30-35 mice with antibodies specific to vGAT. We found that W1989R 480 kDa ankyrin-G is capable of stabilizing Pinceau GABAergic synapses on the soma and AIS of Purkinje neurons (Supplementary Fig. 5A and B). These data show that, while the formation of GABAergic synapses in the cerebellum is mediated by 480 kDa ankyrin-G^16^, it is through a GABARAP-independent mechanism. Overall, these results demonstrate that stabilization of cell surface GABAA receptors mediated by 480 kDa ankyrin-G and GABARAP is a cell type- and brain region-specific mechanism for the proper formation and function of GABAergic synapses *in vivo*.

### Homozygous *Ank3* W1989R mice show altered network synchronization

Parvalbumin-positive (PV+) GABAergic interneurons are critical for the synchronization of forebrain networks due to their rhythmic, fast-spiking electrophysiological properties, and because a single PV+ interneuron can synapse onto hundreds or thousands of pyramidal neurons simultaneously^21, 22^. There are two main subtypes of PV+ interneurons: the PV+ basket cells, which synapse onto the soma and proximal dendrites of pyramidal neurons, and PV+ chandelier cells, which innervate the AIS. Decreased GABAergic synapses and sIPSC frequency observed in the *Ank3* W1989R mouse model (Fig. 2 and Supplementary Fig. 3) suggested a reduction in presynaptic connectivity or reduced density of PV+ interneurons. To test this hypothesis, we measured PV+ cell number and function in layer II/III of the somatosensory cortex. Immunostaining of coronal brain sections with anti-PV antibodies revealed no detectable changes in the density of PV+ interneurons in *Ank3* W1989R mice compared to WT (Fig. 3A and B). To evaluate the firing properties of PV+ interneurons, we evoked AP firing by injecting somatic current in acute brain slices of P25-48 homozygous *Ank3* W1989R or WT mice. *Ank3* W1989R PV+ interneurons maintained their fast-spiking electrophysiological properties and demonstrated similar AP frequency and amplitude compared to WT PV+ neurons (Fig. 3C, E, and F). We observed a significant increase in the membrane resistance in *Ank3* W1989R PV+ interneurons compared to WT (Fig. 3D), suggesting decreased ion channel expression at the plasma membrane.

PV+ interneurons comprise approximately 40% of the total number of interneurons in the somatosensory cortex^23^. To examine the effect of the *Ank3* W1989R mutation on two additional subclasses of GABAergic interneurons, we performed wholecell patch clamp recording on regular spiking non-pyramidal interneurons (RSNP) and irregular spiking interneurons (IS). Despite a large decrease in detectable GABAergic synapse connectivity, there was no significant difference in action potential frequency or amplitude of RSNP interneurons or IS interneurons in *Ank3* W1989R mice compared to WT (Supplementary Table 2). Moreover, there was significant reduction in AP threshold from RSNP and IS interneurons between WT and *Ank3* W1989R mice; however, the input resistance, resting membrane potential, AP amplitude, and AP *τ* were unchanged (Supplementary Table 2). These data suggest that loss-of-function of the 480 kDa isoform of ankyrin-G results in decreases in GABAergic synaptic connectivity; however, GABAergic inhibitory interneurons are present at normal density and maintain function.

PV+ interneurons are responsible for the generation of gamma oscillations, which reflect the precise synchronization of local neuronal networks^22^. We hypothesized that, because *Ank3* W1989R mice exhibit reductions in somatodendritic and AIS GABAergic synapses, network synchronization would be decreased. To evaluate the synchronization of neuronal ensembles, we used planar multi-electrode arrays to record kainate-induced gamma oscillations in acute slices of hippocampus from WT and homozygous *Ank3* W1989R mice^24^ (Fig. 4A and B). The power of the kainate-induced gamma oscillations was ~30% decreased in CA1 and CA3 hippocampal regions in *Ank3* W1989R mice compared to WT (Fig. 4C, D, E, and F). This significant reduction in gamma oscillations in *Ank3* W1989R mouse hippocampus suggests disruptions in network synchrony, consistent with reduced connectivity of PV+ interneurons.

**Fig. 4:**
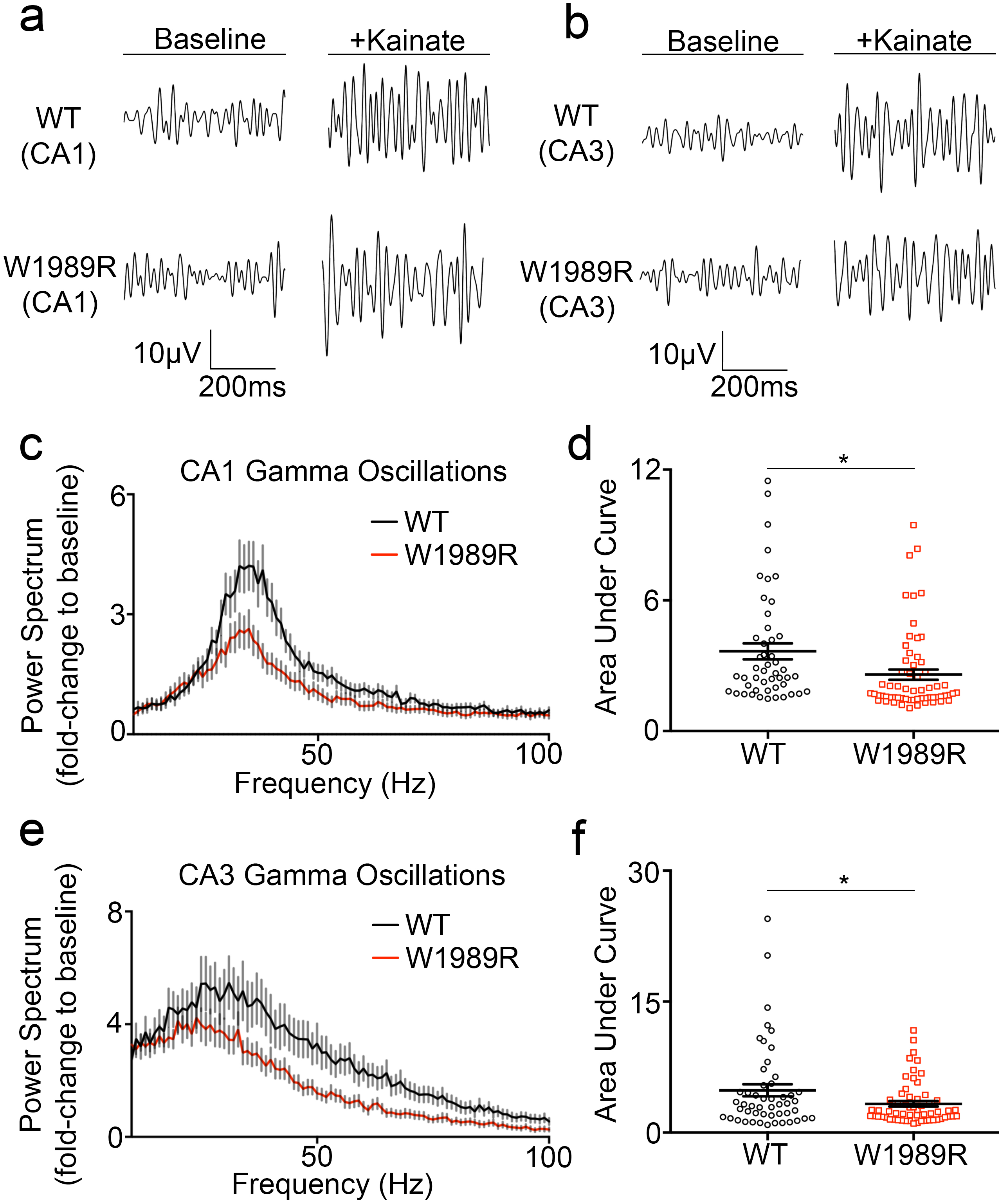
Reduced gamma oscillations in the hippocampus of *Ank3* W1989R mice. **(a)** Representative traces of kainate-induced gamma oscillations from local field potential (LFP) recordings in CA1 hippocampal neurons and **(b)** CA3 hippocampal neurons of WT (top) or *Ank3* W1989R homozygous (bottom) mice in acute brain slices. **(c)** Power spectral analysis from CA1 hippocampus in acute brain slices of P24-40 WT (black circles) and *Ank3* W1989R homozygous (red squares) mice plotted as fold-change to baseline signal. **(d)** Quantification of the area under the curve for gamma band (30-60Hz) from WT (black circles) and *Ank3* W1989R homozygous (red squares). *t-test* *P = 0.0118 (WT: 3.66 ± 0.4, N=3, n=48; W1989R: 2.59 ± 0.2, N=3, n=62). **(e)** Power spectral analysis from CA3 hippocampus WT (black circles) and *Ank3* W1989R homozygous (red squares) mice. **(f)** Quantification of the area under the curve for gamma band (30-60Hz) from WT (black circles) and *Ank3* W1989R homozygous (red squares). *t-test* *P=0.0343 (WT: 4.85 ± 0.7, N=5, n=50; W1989R: 3.3 ± 0.3, N=5, n=59).

### Dendritic spine density and function are reduced in *Ank3* W1989R neurons

GABAergic interneurons control the excitability of glutamatergic pyramidal neurons by modulating their spike timing and firing rate. The loss of inhibitory tone, similar to that observed here in *Ank3* W1989R mice, has been linked to pathological neuronal hyperexcitability^25^. To address the effect of decreases in GABAergic synapses on AP firing rates, we compared evoked APs in acute brain slices of P25-48 homozygous *Ank3* W1989R and WT mice. The frequency of AP firing was significantly increased in *Ank3* W1989R cortical and CA1 hippocampal neurons compared to WT (Fig. 5A, C, D, F). Moreover, the maximum firing rate per neuron was two-fold higher in *Ank3* W1989R neurons versus WT (Fig. 5B and E). These data demonstrate that cortical and hippocampal pyramidal neurons are hyperexcitable *in vivo* following 480 kDa ankyrin-G loss-of-function at GABAergic synapses.

**Fig. 5:**
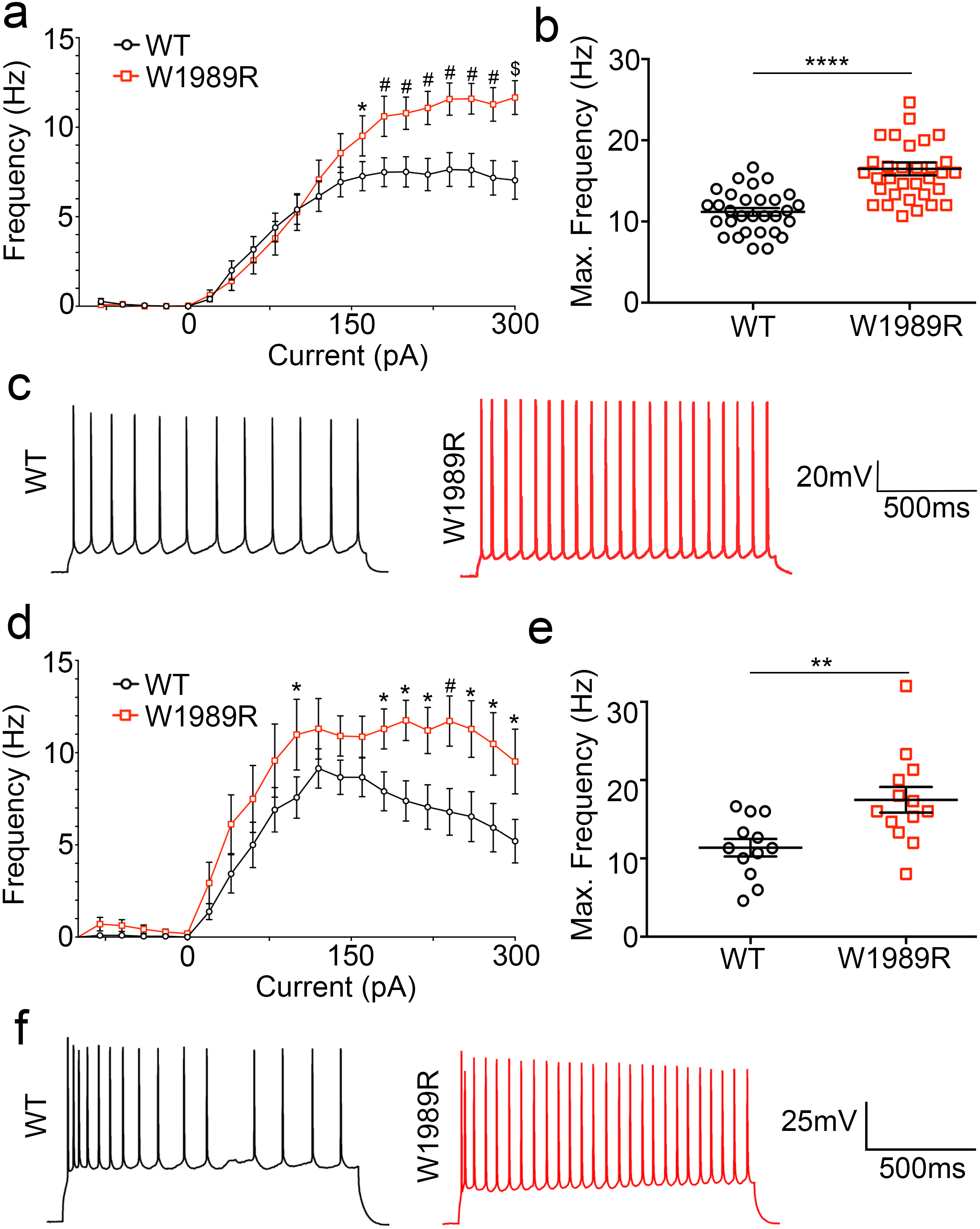
Increased firing rate of *Ank3* W1989R cortical and CA1 hippocampal pyramidal neurons. **(a)** Evoked AP frequency from cortical neurons for WT (black circles) or *Ank3* W1989R homozygous (red squares) in acute brain slices. *Two-way ANOVA, Tukey’s post hoc* *P < 0.05, ^#^P <0.001, ^$^P <0.001 (WT: n=40; W1989R: n=40). **(b)** Quantification of average maximum frequency of pyramidal neurons in layer II/III somatosensory cortex in WT (black circles) and *Ank3* W1989R homozygous (red squares) mice. *t-test* ****P < 0.0001 (WT: 11.18 ± 0.5, n=30; W1989R: 16.49 ± 0.8, n=33). **(c)** Representative AP traces from cortical pyramidal neurons of WT (left, black) and *Ank3* W1989R homozygous (right, red) mice. Scale bar: 500 ms. **(d)** Evoked AP frequency from CA1 hippocampal neurons for WT (black circles) or *Ank3* W1989R homozygous (red squares) in acute brain slices. *Two-way ANOVA, Tukey’s post hoc* *P < 0.05, ^#^P <0.001 (WT: n=27; W1989R: n=27). **(e)** Quantification of average maximum frequency of pyramidal neurons in CA1 hippocampus of WT (black circles) and *Ank3* W1989R homozygous (red squares) mice. *t-test* **P = 0.0061 (WT: 11.38 ± 1.1, n=12; W1989R: 17.49 ± 1.6, n=13). **(f)** Representative traces from CA1 hippocampal pyramidal neurons of WT (left) and *Ank3* W1989R homozygous (right) mice. Scale bar: 500 ms. Data shown as mean ± SEM.

Neurons have developed multiple intrinsic mechanisms to maintain excitability homeostasis and stabilize network activity. The AIS has been proposed to participate in activity-dependent plasticity and decreasing AIS length may be one mechanism to reduce excitability. We measured AIS length in cortical brain sections and in dissociated hippocampal neurons from WT and mutant mice. We found that AIS length was approximately 30% shorter in *Ank3* W1989R mice compared to WT (Supplementary Fig. 6). Another neuronal mechanism to compensate for the lack of inhibitory tone is to decrease AMPA receptor surface expression and decrease dendritic spine density ^26^. We therefore examined dendritic spine density in dissociated hippocampal neurons from *Ank3* W1989R mice and observed a significant decrease in the total number of dendritic spines compared to WT (Fig. 6A and B). Consistent with this result, whole-cell patch clamp recordings from CA1 hippocampal neurons in brain slices showed significant decreases in mEPSC amplitude in *Ank3* W1989R mice compared to WT, suggesting a reduction in dendritic AMPARs (Fig. 6C, D, and E). Moreover, mEPSC frequency was increased in *Ank3* W1989R neurons (Fig. 6D). A previous study showed that 190 kDa ankyrin-G plays a critical role in modulation of dendritic spine morphology and AMPA receptor postsynaptic stability^27^. Western blot analysis of cortical lysates from *Ank3* W1989R mice showed a 50% decrease in 190 kDa ankyrin-G expression, with no change in the expression of the 270- or 480 kDa splice variants, compared to WT lysates (Fig. 6F). Overall, these data suggest a novel intrinsic mechanism to compensate for neuronal hyperexcitability by decreasing the expression of 190 kDa ankyrin-G and subsequently dendritic spine density and function in neurons with reduced inhibitory tone.

**Fig. 6:**
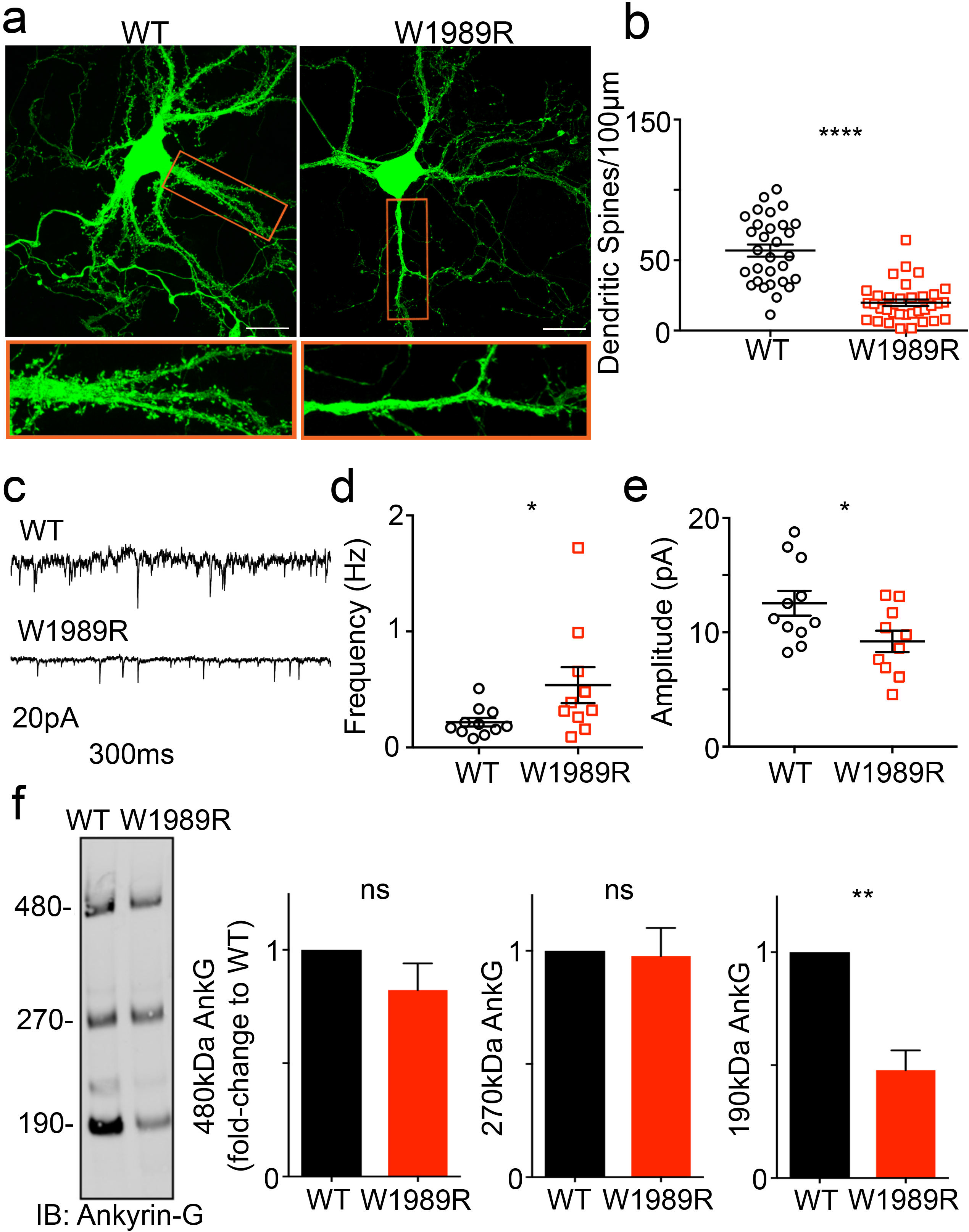
Decreased spine density and function and reduced 190 kDa ankyrin-G expression levels in *Ank3* W1989R neurons. **(a)** Representative images of dissociated hippocampal cultured neurons from WT (left) and *Ank3* W1989R homozygous (right) mice at 21DIV filled with soluble eGFP. Scale bar: 20 μm. **(b)** Quantification of the total number of dendritic spines per 100 μm per neuron. *t-test* ****P < 0.0001 (WT: 56.91 ± 4.3, N=3, n=30; W1989R: 19.78 ± 2.3, N=3, n=34). **(c)** Representative traces of spontaneous excitatory post-synaptic currents (sEPSCs) from CA1 hippocampal neurons in WT (top) and *Ank3* W1989R (bottom) slices. Scale bar: 300 ms. **(d)** Quantification of mEPSC frequency in WT (black circles) and *Ank3* W1989R (red squares) brain slices. *t-test* *P = 0.049 (WT: 0.22 ± 0.04 Hz, n=11; W1989R: 0.54 ± 0.15 Hz, n=10). **(e)** Quantification of mEPSC amplitude in WT (black circles) and *Ank3* W1989R (red squares) brain slices. *t-test* *P = 0.032 (WT: 12.5 ± 1.1 pA, n=11; W1989R: 9.21 ± 0.9 pA, n=10). **(f)** Western blot analysis from cortical lysates of P30 WT (left) and *Ank3* W1989R homozygous (right) mice. Blots were probed with antibodies to total ankyrin-G. Quantification of relative expression levels of 480 kDa ankyrin-G *t-test* P = 0.21 (WT: 1.0 ± 0, n=3; W1989R: 0.82 ± 0.11, n=3), 270 kDa ankyrin-G *t-test* P = 0.86 (WT: 1.0 ± 0, n=3; W1989R: 0.98 ± 0.12, n=3), and 190 kDa ankyrin-G *t-test* **P = 0.004 (WT: 1 ± 0 n=3; W1989R: 0.48 ± 0.09, n=3). Data normalized to WT controls. Data shown as mean ± SEM.

### Identification of *ANK3* W1989R in a family with BD

Disruptions in forebrain circuitry and network-level activity observed in *Ank3* W1989R mice predict that patients expressing this variant may experience altered brain activity and mood-related behaviors. According to the most recent data from the gnomAD project, the ANK3 W1989R variant 10:61834674 (GRCh37.p13) is found in approximately 1: 10,000 European Americans^28^. We used whole genome and exome sequencing on blood samples obtained through the Heinz C. Prechter Bipolar Research Program at the University of Michigan to identify a patient expressing the ANK3 W1989R variant (Fig. 7A). We confirmed the presence of the variant by extracting DNA from fibroblasts derived from the proband and performing nested PCR followed by Sanger sequencing. The proband (II:2, age 45) was diagnosed with BD type I characterized by recurrent mania and depression with an age of onset of 17 years, with current successful maintenance on lithium (1200 mg daily) and a benzodiazepine (pro re nata) PRN at bedtime. The proband had a brief (<3 months) exposure to antipsychotic medication (chlorpromazine), but no history of treatment with antidepressant medication. To determine whether other family members also carried the ANK3 W1989R variant, we expanded our studies to include the proband’s parents (I:1, I:2) and sister (II:1). We extracted DNA from whole blood and performed nested PCR of the region flanking ANK3 W1989 followed by Sanger sequencing. The proband (II:2), the mother (I:2) and the sister (II:1) were heterozygous for the ANK3 W1989R variant (Fig. 7A and B). The mother (age 73) was diagnosed with BD type I with age of onset in her mid-30’s. She is currently treated in the community with lamotrigine (50 mg daily), clomipramine (50 mg daily) and lorazepam (1-3 mg daily), as needed, for anxiety. The proband’s sister (age 50) was diagnosed with BD type II, with a prepubertal onset of mood instabilities and multiple episodes of depression and hypomania with mixed affective features. Her treatment currently includes antidepressants (fluoxetine 60 mg and buproprion 300 mg daily), an anticonvulsant (lamotrigine 400 mg daily), a stimulant (amphetamine 60 mg daily), and a hypnotic (temazepam 30 mg at bedtime). The proband’s daughter (age 19) was diagnosed with major depression, but no DNA sample was available. The father (I:1) is WT for W1989 and has no history of depression or treatment of any psychiatric disorder (Fig. 6B). This is the first reported characterization of the *ANK3* W1989R human variant 10:61834674 tgg>cgg (Fig. 6C). Similar to results from *Ank3* W1989R mutant mice, patients carrying the *ANK3* W1989R variant have survived into adulthood, demonstrating that this variant is compatible with normal lifespan in humans. However, the presence of the *ANK3* W1989R variant in three affected patients is consistent with a potential effect of this variant on neuronal activity and mood-related behaviors.

**Fig. 7:**
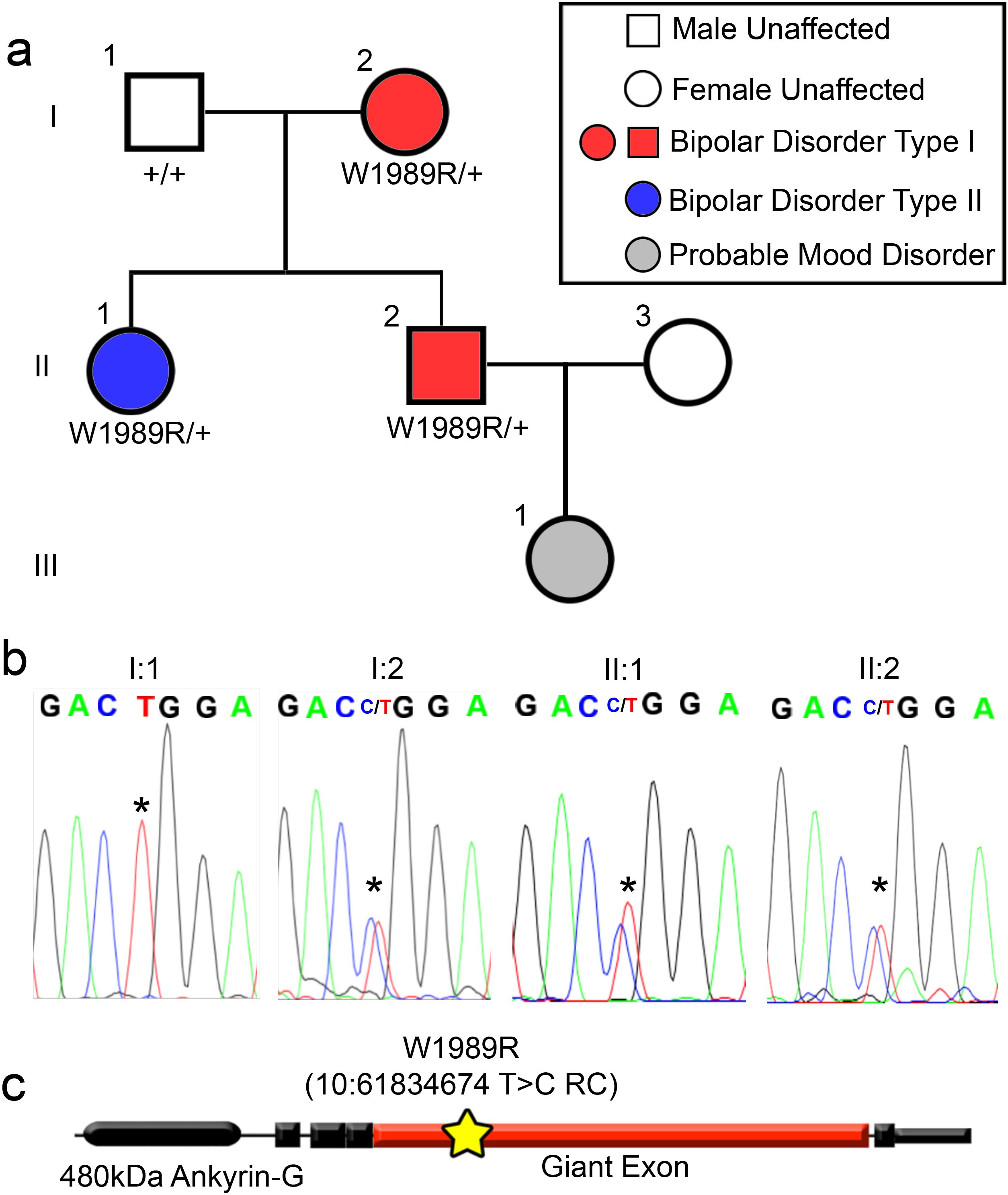
*ANK3* W1989R variant identified in a family with bipolar disorder. **(a)** Pedigree of individuals heterozygous for *ANK3* W1989R mutation diagnosed with BD. Red: diagnosis of BD type I. Blue diagnosis of BD type II. Gray: individuals with signs and symptoms of mood disorders. White: neurotypical individuals. Circles: female. Squares: male. The affected proband is individual II:2 represented by a red square. **(b)** Chromatogram from Sanger sequencing confirming presence of *ANK3* W1989R heterozygous mutation in affected mother I:2, affected proband II:2, and proband’s affected sister II:1. Asterisk: site of missense mutation in affected individuals I:2, II:1, and II:2 or wild-type allele in I:1. **(c)** Schematic of the 480 kDa splice variant of ankyrin-G. Yellow star: site of W1989R variant within the giant exon (red) of ankyrin-G. Genomic position: 10:6183467; Variant: T > C. RC: reverse complement.

## DISCUSSION

Neuropsychiatric diseases, such as BD and schizophrenia, are highly heritable. Significant progress has been made in the past decade identifying genetic risk factors associated with neuropsychiatric diseases including common single nucleotide polymorphisms (SNPs) through GWAS, copy number variants, and rare inherited and *de novo* variants^29^. For many neurological diseases, common genetic variants have minimal impact on disease susceptibility, thus it is important to understand how rare genetic variants impact brain function and underlie the pathophysiology of neuropsychiatric disease. *ANK3* is among the most consistent and significant genes associated with BD^30–33^; however, the mechanisms by which *ANK3* variants contribute to pathophysiology are not known. Although *ANK3* has been associated with multiple neurological disorders, the cellular and molecular mechanisms by which its loss-of-function contributes specifically to BD are poorly understood.

In this study, we generated homozygous *Ank3* W1989R knock-in mice to investigate ankyrin-G/GABARAP interactions and to understand the effects of 480 kDa ankyrin-G loss-of-function at GABAergic synapses *in vivo*. We found that *Ank3* W1989R mice have reduced GABAergic synapses on the AIS and somatodendritic domain of cortical and hippocampal pyramidal neurons, while the density and function of PV+ GABAergic interneurons are maintained. Gamma oscillations were reduced in *Ank3* W1989R hippocampus, suggesting disruptions in network synchrony, consistent with reduced connectivity of PV+ interneurons. We found that AIS length and dendritic spine density and function were reduced in *Ank3* W1989R cortical and hippocampal neurons, suggesting that regulation of ankyrin-G-dependent domains may be a neuronal mechanism of homeostasis. Finally, we identified the *ANK3* W1989R variant (rs372922084) in a family with BD, suggesting that *ANK3* W1989 is a critical residue involved in disease mechanisms in human patients.

The decrease in AIS length in *Ank3* W1989R mice may be due to neural plasticity changes, as recent reports have shown that alterations in AIS length, location, and/or ion channel surface expression may occur following fluctuations in neuronal activity in an attempt to maintain homeostasis of intrinsic excitability^34^. Further, we propose that the decreased dendritic spine density and function observed in *Ank3* W1989R mice may be an additional compensatory mechanism to modulate neuronal hyperexcitability. The W1989R mutation is specific to the giant exon, only included in the 270 kDa and 480 kDa splice variants of ankyrin-G, which suggests that the observed decrease in 190 kDa ankyrin-G expression may be a compensatory effect to maintain excitatory/inhibitory balance. Further, the observed increase in mEPSC frequency may be explained by hyperexcitability of upstream glutamatergic neurons due to the decrease in inhibitory synapses. Our work shows that the interaction between 480 kDa ankyrin-G and GABARAP is necessary for stabilizing forebrain GABAergic synapses, while additional mechanisms are involved in the formation of GABAergic synapses in the cerebellum. Previous studies have shown that deletion of 480 kDa ankyrin-G *in vivo* results in the loss of GABAergic synapses at the AIS of cerebellar Purkinje neurons ^13, 16^. Further, a study by Ango et al. showed an ankyrin-G-dependent subcellular gradient of neurofascin-186 at the AIS of Purkinje neurons is necessary for organizing GABAergic synapses ^35^. Thus, ankyrin-G plays a critical role in stabilizing GABAergic synapses thorough unique brain region- and cell type-specific mechanisms.

Several mouse models have been generated to understand how *Ank3* contributes to neuronal development and function. Specific deletion of the exon encoding the giant 270- and 480 kDa isoforms of ankyrin-G results in the loss of all known AIS components, gross malformations in the morphology and total number of nodes of Ranvier, and decreased GABAergic inhibitory synapses. This genetic manipulation also results in lethality at P20, preventing the study of mature animals^16^. A recent study generated a mouse model with forebrain-specific deletion of all splice variants of ankyrin-G^36^. These ankyrin-G conditional null mice demonstrate “mania-like” behaviors including hyperactivity, increased anxiety, and exploratory behavior, as well as depressive-like behaviors following social defeat stress^36^. The behavioral phenotypes identified in this model were rescued following administration of lithium and valproic acid^36^. Although the animals survived well into adulthood, the loss of multiple cellular domains made it difficult to determine which ankyrin-G-dependent domain underlies the observed behavioral phenotypes related to BD. A study by Lopez et al. showed that deletion of *Ank3* exon 1b, which affects ankyrin-G splice variants encoded by alternative first exon and N-termini splicing, results in decreased ankyrin-G expression at the AIS of PV+ interneurons^37^. These investigators found that gene dosage-dependent reductions in expression of ankyrin-G corresponded to disease severity. Heterozygous mice displayed behavioral phenotypes representative of BD, whereas homozygous mice demonstrated epilepsy and sudden death^37^. Another study compared the behaviors of *Ank3* heterozygous null mice with mice in which ankyrin-G was deleted specifically in the hippocampal dentate gyrus using short hairpin RNA (shRNA) viral vectors^38^. Both models demonstrated reduced anxiety and increased impulsivity, behavioral phenotypes related to BD^38^. These behaviors were reversed following chronic lithium treatment^38^. Taken together, these studies, along with the present work, provide evidence that *ANK3* is critical for normal brain function.

Abnormalities in excitatory: inhibitory circuit balance have been implicated in the pathophysiology of BD and schizophrenia ^39–45^. Postmortem brains of BD patients revealed decreased expression of the GABAergic synapse marker GAD67, the GABA transporter GAT1, and various GABAA-receptor subunits^3, 4, 46^. We found morphological and functional reductions in cortical and hippocampal GABAergic synapses in our mouse model, consistent with GABAergic dysfunction reported in patients with BD^39^. Cortex and hippocampus are key brain regions associated with the cognitive, emotional, and mood-related behaviors characteristic of BD^1^. Further, abnormalities in dendritic spines have been associated with neuropsychiatric diseases including BD, schizophrenia, autism spectrum disorders, and intellectual disability^40, 41 43–45^. We found reduced density and function of dendritic spines in our mouse model and proposed that these effects act to reduce hyperexcitability and compensate for the lack of inhibitory input. Previous studies have observed impairments in gamma oscillations in patients with neuropsychiatric disease, including BD and schizophrenia^2, 6–8, 25^. Further, recent reports have suggested that gamma oscillations may serve as biomarkers for diagnosing and tracking treatment response in individuals with BD^5^. Consistent with these studies, we found significant decreases in hippocampal gamma oscillations in the *Ank3* W1989R mice. Thus, our cellular and functional characterization of inhibitory and excitatory synaptic dysfunction in the *Ank3* W1989R mouse model may inform the future development of novel therapeutics for the treatment of BD and other neurological diseases involving altered excitatory: inhibitory balance.

*ANK3* variants have been associated with schizophrenia^47^, autism^48^, epilepsy^37^, and intellectual disability in addition to BD^49^. Variants in *ANK3* may contribute to these neurological diseases by affecting ankyrin-G expression levels, disrupting protein folding, impacting specific splice variants, or preventing ankyrin-G from interacting with critical binding partners, as observed for GABARAP in the *Ank3* W1989R mouse model. Thus, it remains important to understand how disease-associated variants in *ANK3* affect ankyrin-G function and contribute to disease pathology. Although *ANK3* W1989R is a rare variant, the data reported in this work may have broader impacts on BD patients carrying other variants that reduce ankyrin-G expression. Several independent GWAS studies revealed *ANK3* BD-associated SNPs near the 5’ non-coding region, which could potentially lead to altered expression levels of different isoforms of ankyrin-G^33, 50–52^. Moreover, studies using postmortem brains from BD patients found reduced *ANK3* mRNA expression^53, 54^. Consistent with the *ANK3* W1989R variant, BD-associated rare variants have been detected within alternatively spliced exons of ankyrin-G^55^. One potential explanation to describe the genetic etiology of BD is that rare variants in *ANK3* have a high penetrance due to dysfunction of the gene or encoded splice variant. Alternatively, multiple common variants with low penetrance may lead to BD due to the polygenic nature of disease inheritance. In support of this hypothesis, rare variants in several genes involved in GABAergic and glutamatergic neurotransmission as well as voltage-gated calcium channels contribute to increased risk of BD, as they may result in similar endophenotypes as individuals with *ANK3* mutations^56^. Ultimately, it will be important to continue to evaluate how patient-specific rare variants affect the expression and function of the different splice variants of ankyrin-G to provide insight on additional pathways that may contribute to BD and other neurological diseases.

## MATERIALS AND METHODS

### Constructs, Protein Expression, and Purification

The coding sequence of the GABARAP (UniProt: Q9DCD6) construct was PCR amplified from a mouse brain cDNA library. The coding sequence of ankyrin-G construct was PCR amplified from the full-length rat 270 kDa ankyrin-G (UniProt: O70511) template. All point mutations were generated using the Quikchange II XL site-directed mutagenesis kit and confirmed by DNA sequencing. All of the constructs used for protein expression were cloned into a home-modified pET32a vector. Recombinant proteins were expressed in BL21 (DE3) *E. coli* cells with induction of 0.25 mM IPTG at 16°C. The N-terminal Trx-His_6_-tagged proteins were purified using Ni^2+^-NTA agarose affinity columns followed by size-exclusion chromatography (Superdex 200 column from GE Healthcare) in the final buffer containing 50 mM Tris-HCl, 1 mM DTT, and 1 mM EDTA, pH 7.8 with 100 mM NaCl. The Trx-His_6_ tag was removed by incubation with HRV 3C protease and separated by size exclusion columns or reverse usage of Ni^2+^-NTA columns when needed.

### Crystallography

Crystallization of the ankyrin-G/GABARAP complex was performed using the sitting drop vapor diffusion method at 16 °C. Crystals of ankyrin-G/GABARAP were grown in solution containing 2.0 M ammonium citrate tribasic and 0.1 M BIS-TRIS propane buffer (pH 7.0). Crystals were soaked in crystallization solution with higher salt concentration (3 M ammonium citrate) for dehydration and cryoprotection. All datasets were collected at the Shanghai Synchrotron Radiation Facility BL17U1 or BL19U1 beamline at 100 K. Data were processed and scaled using HKL2000 or HKL3000. Structure was solved by molecular replacement using PHASER with apo form structure of GABARAP (PDB: 1KJT) as the searching model. The ankyrin-G peptide was manually built according to the F_o_-F_c_ difference maps in COOT. Further manual model adjustment and refinement were completed iteratively using COOT and PHENIX. The final model was validated by MolProbity and statistics are summarized in Table 1. All structure figures were prepared by PyMOL (http://www.pymol.org).

### Isothermal Titration Calorimetry Assay

Isothermal titration calorimetry (ITC) measurements were carried out on a VP-ITC Microcal calorimeter at 25 °C with the titration buffer containing 50 mM Tris-HCl, pH 7.8, 100 mM NaCl, 1 mM DTT, and 1 mM EDTA. For a typical experiment, each titration point was performed by injecting a 10 μL aliquot of protein sample (200 μM) into the cell containing another reactant (20 μM) at a time interval of 120 seconds to ensure that the titration peaks returned to the baseline. 27 aliquots were titrated in each individual experiment. The titration data were analyzed using the program Origin 7.0 and fitted by the one-site binding model.

### Generation of W1989R Mouse Model

A knock-in mouse was generated by inserting the tryptophan to arginine mutation corresponding to human W1989R within the neuronal-specific giant exon of the mouse *Ank3* gene, which corresponds to exon 37 of human *ANK3*, ENST00000280772. The exon 37 plasmid contained the W1989R mutation and a neomycin resistance cassette. The neomycin resistance cassette was flanked by LoxP sites containing flippase recognition target (FRT) sites. The linearized construct was introduced into 129S6/SvEvTac-derived TL1 embryonic stem (ES) cells by electroporation. ES cells selected for the W1989R mutation using neomycin were injected into C57BL/6NHsd blastocysts. High percentage chimeric animals were obtained and bred to C57BL/6NHsd mice to produce heterozygous animals. The neo cassette was excised by crossing with the W1989R *Ank3* mouse containing a floxed neo cassette with a Sox2-Cre mouse [B6.Cg-Tg(Sox-cre)1Amc/J, stock number 008454; The Jackson Laboratory]. Mutant mice were backcrossed for at least six generations to C57BL6/J mice from the Jackson Laboratory and were compared to C57BL/6J mice as WT controls. All mouse production was provided by the Duke Cancer Institute Transgenic Mouse Facility. All experiments were performed in accordance with the guidelines for animal care of the Institutional Animal Care and Use Committee (IACUC) and University Laboratory Animal Management (ULAM) at the University of Michigan.

### Immunocytochemistry of Brain Sections

For immunohistochemistry, P30-35 mice were administered a ketamine/xylazine mixture (80 mg/kg body weight ketamine and 10 mg/kg xylazine) via intraperitoneal injection. The mice were sacrificed by cardiac perfusion of PBS followed by 4% paraformaldehyde and the brain was immediately removed and fixed overnight in 4% paraformaldehyde. The next day, the brains were processed using a standard single-day paraffin preparation protocol (PBS wash followed by dehydrations through 70, 95, and 100% ethanol with final incubations in xylene and hot paraffin under vacuum) using a Leica ASP 300 paraffin tissue processor. Paraffin sections were cut 7 μm thick using a Leica RM2155 microtome and placed on glass slides. Sections were deparaffinized and rehydrated using a standard protocol of washes: 3 × 4-min xylene washes, 2 × 2-min 100% ethanol washes, and 1 × 2-min 95%, 80%, and 70% ethanol washes followed by at least 5 min in ddH_2_O. Antigen retrieval was then conducted by microwaving the deparaffinized brain sections for 20 min in 10 μM sodium citrate. Sections were cooled, washed for 15 min in ddH_2_O, rinsed in PBS for 5 min, and blocked using blocking buffer (5% BSA, 0.2% Tween 20 in PBS) for 1 hour at room temperature. Slides were incubated overnight at 4°C with primary antibodies diluted in blocking buffer. On the following day, slides were washed 3 times for 15 min with PBS containing 0.2% Tween 20, incubated with secondary antibodies diluted in blocking buffer for 1 hour at room temperature, washed 3 times for 15 min, and mounted with Prolong Gold.

### Neuronal Culture and Transfection

Hippocampi were dissected from postnatal day 0 (P0) mice, treated with 0.25% trypsin and 100 μg/ml DNase in 2 mL HBSS with 10 mM HEPES, and then gently triturated through a glass pipette with a fire-polished tip. The dissociated neurons were then plated on poly-D-lysine and laminin-coated 35 mm MatTek dishes in 0.5 mL of Neurobasal-A medium containing 10% (vol/vol) FBS, B27 supplement, 2 mM glutamine, and penicillin/streptomycin. On the following day, 2.5 mL of fresh Neurobasal-A medium containing 1% FBS, B27, glutamine, and penicillin/streptomycin was added to the dish. AraC was added at 1:1000 to protect against glial and fibroblast overgrowth. Plates were returned to incubation at 37°C until experimentation. To fill the cells with soluble GFP, the dissociated hippocampal cultures were transfected with eGFPN1 plasmid. Briefly, 1 μg eGFPN1 plasmid was added to 100 μl of Neurobasal-A and, in a second tube, 3 μl of Lipofectamine 2000 was added to 100 μL of Neurobasal-A. The two tubes were mixed and incubated for 15 min at room temperature. The neuronal growth media was then aspirated from the dishes and saved, the transfection was added dropwise to 14 DIV neurons, and the transfected cells were incubated at 37°C for 1 hr. The transfection mixture was aspirated and the original neuronal growth media was added. The cells were maintained in culture until 21 DIV and fixed for immunofluorescence as described below.

### Immunofluorescence of Cultured Neurons

Dissociated hippocampal neurons were fixed for 15 min at room temperature with 4% paraformaldehyde, followed by methanol for 10 min at −20°C, and blocked with blocking buffer (5% BSA, 0.2% Tween 20 in PBS). Primary antibodies were diluted in blocking buffer and incubated at 4°C overnight. The following day, cells were washed 3 × 15 min with PBS containing 0.2% Tween 20, incubated with secondary antibodies diluted in blocking buffer for 1 hour at room temperature, washed 3 × 15 min, and mounted with Prolong Gold.

### Confocal Microscopy

Samples were imaged on a Zeiss LSM 880 with a 60X NA1.4 Oil/DIC Plan-Apochromat objective and excitation was accomplished using 405, 488, 561, and 633 nm lasers.

### In Vitro Electrophysiology Recordings and Analysis

Brains were obtained from WT C57BL/6J or *Ank3* W1989R mutant mice between P25-48. The animals were decapitated under isoflurane and USP anesthesia, the brain was then quickly removed from the skull and placed in 4°C slicing solution containing 62.5 mM NaCl, 2.5 mM KCl, 1.25 mM KH_2_PO_4_, 26 mM NaHCO_3_, 5 mM MgCl_2_, 0.5 mM CaCl_2_, 20 mM glucose and 100 mM sucrose (pH maintained at 7.4 by saturation with O_2_/CO_2_, 95/5% respectively). Coronal brain slices (300-350 μm thick) containing layers II/III somatosensory cortex and hippocampus were cut with a microtome (VF-300, Compresstome). The slices were then transferred to a holding chamber and maintained at room temperature in artificial cerebrospinal fluid (ACSF) containing 125 mM NaCl, 2.5 mM KCl, 1.25 mM KH_2_PO_4_, 26 mM NaHCO_3_, 1 mM MgCl_2_, 2 mM CaCl_2_ and 20 mM glucose, pH 7.4 (with 95%O_2_ and 5%CO_2_ bubbling through the solution) for at least 1 hour prior to recording. After equilibration, individual slices were transferred to the recording chamber continuously perfused with ACSF (1-2 mL/min). Recording micropipettes were pulled from borosilicate glass capillaries (1.5 mm O.D. HARVARD APPARATUS) for a final resistance of 3-6 MΩ and filled with a solution containing 135 mM K-Gluconate, 4 mM NaCl, 0.4 mM GTP, 2 mM Mg-ATP,0.5mM CaCl2, 5 mM EGTA and 10 mM HEPES. The signals were recorded with an Axoclamp 700B amplifier (Axon Instruments, Union City, CA), low pass filtered at 10 kHz. Current clamp recordings were obtained from neurons in layers II/III of somatosensory cortex; the cells in this area were identified using a Nikon Eclipse FN-1 microscope with a 40X water-immersion objective and a DAGE-MTI IR-1000 video camera. Whole-cell patch-clamp recordings with a high cell resistance (greater than 8 GΩ before break-in) were obtained for cells according to the availability. The neurons were characterized electrophysiologically by applying negative and positive current pulses of 20 pA and 1500 ms to calculate the maximum frequency and positive pulses of 50 ms to measure the features for the single AP. For sIPSC and mIPSC recordings in voltage-clamp configuration, the K-gluconate in the internal solution was replaced by CsCl and the recordings were acquired at 2 kHz fixing the voltage at −80 mV. The IPSCs were recorded in the presence of the N-methyl-D-aspartate receptor blockers and non-N-methyl-D-aspartate glutamate receptors, AP-5 (50 mM) and CNQX (10 μM). For measurement of mIPSCs, 1 μM tetrodotoxin (TTX) was added to the superfusion to block synaptic responses dependent on the AP.

Access resistance was monitored throughout the experiment and experiments were canceled if changes greater than 20% occurred. The mEPSCs were measured in the presence of bicuculline (10 μM) and TTX (1 μM) while holding the resting membrane potential at −70 mV using K-gluconate internal solution. The peak events were searched automatically using Minianalysis (Synaptosoft Inc.) and visually monitored to exclude erroneous noise. Both the frequency and amplitude of the events and their distribution were analyzed. Mean values were compared using the student’s ŕ-test. All data are presented as mean ± SEM.

### *In Vitro* Kainate-Induced Oscillations

WT C57BL/6J or *Ank3* W1989R mice between P25-40 were anesthetized with isoflurane. Mice were perfused intracardially with ice cold (4°C) modified N-methy-D-glucamine (NMDG) HEPES artificial cerebrospinal fluid (aCSF) consisting of: 93 mM NMDG 2.5 mM KCl, 0.5 mM CaCl_2_, 10 mM MgCl_2_, 1.2 mM NaH_2_PO_4_, 20 mM HEPES, 25 mM dextrose, 5 mM ascorbic acid, 2mM thiourea and 3 mM Na-pyruvate. pH was maintained at 7.4 by saturation with O_2_/CO_2_, (95/5%, respectively). Horizontal hippocampal sections (300 μm thick) were prepared with a Leica VT1200 vibratome. Sections were bi-laterally hemisected and transferred to a holding chamber maintained at 33° C for 10-12 min, and then transferred to a holding chamber with aCSF consisting of: 126mM NaCl, 3mM KCl, 2mM CaCl_2_, 1 mM MgCl_2_, 1.25 mM NaH_2_PO_4_ 25 mM NaHCO_3_ and 10 mM dextrose. pH was maintained at 7.4 by saturating aCSF with 95%O_2_/5% CO_2_ at 33° C for 35min. Sections were then transferred to room temperature for 15 min and were mounted on perforated multi-electrode array (pMEAs)

(Multichannel Systems, Reutlingen, Germany). Sections were secured to the surface of the pMEA surface by using a peristaltic perfusion system (PPS2, Multichannel Systems, Reutlingen, Germany) to create a slight vacuum through the perforations. The secured sections remained submerged in aCSF (29 −31° C, 95% O_2_/5CO^2^%) and were superfused at a rate of 5-7 ml/min. Local field potentials were recorded at 20 kHz. Baseline recordings were obtained for 1 hour. Chemically-induced oscillations were evoked by bath application of 400 nM Kainic Acid (KA) for 1 hour. Oscillations were abolished by bath application of the GABA-A receptor antagonist, bicuculline (10 μM). LFP data were imported into MATALAB, downsampled to 10 kHz and low pass filtered at 400 Hz for analysis. Fast Fourier Transformation (FFT) of LFP data was done in 1 sec bins to calculate the power of oscillations in the last 2 min of each bath condition. Change in oscillatory power was defined as (Power_KA_-Power_aCSF_)/Power_aCSF_. Peak frequency and the area under the curve for gamma band (30 - 60 Hz) were calculated using custom written MATLAB scripts.

### Western Blot

Homogenization buffer consisting of 8M urea, 5% SDS, and 5 mM N-ethylmaleimide was heated to 65°C. Whole brains were dissected from P30-35 mice and immediately frozen in liquid nitrogen. Frozen brains were then ground into a powder using a mortar and pestle. The powder was scraped into a 1.5 mL microcentrifuge tube and hand-dounced in 10 volumes/weight of 65°C homogenization buffer (i.e. 1.5 mL for 150 mg powder). The homogenate was incubated at 65°C for 20 min and then mixed 1:1 with 5× PAGE buffer (5% (wt/vol) SDS, 25% (wt/vol) sucrose, 50 mM Tris, pH 8, 5 mM EDTA, bromophenol blue). The lysates were stored at −80°C until use. The samples (10 μL-volume) were separated on a 3.5-17% gradient gel in 1X Tris buffer, pH 7.4 (40 mM Tris, 20 mM NaOAc, and 2 mM NaEDTA) with 0.2% SDS. Transfer to nitrocellulose was performed overnight at 300 mA at 4°C in 0.5X Tris buffer with 0.01% SDS. Membranes were blocked with 5% non-fat dry milk in TBS and incubated overnight at 4°C with primary antibodies (rabbit total ankyrin-G 1:5,000) diluted in blocking buffer. Membranes were washed 3 × 15 min with TBS-T and incubated for 1 hour with LiCor fluorescent secondaries (1:50,000) in blocking buffer. Membrane were then washed 3 × 15 min in TBS-T, 1 × 5 min TBS, and 1 × 5 min in ddH_2_O before being imaged on LiCor Odyssey Clx imager.

### Antibodies and Reagents

The following antibodies and dilutions were used: rabbit anti-βIV spectrin (1:1000, lab-generated^16^), rabbit anti-total ankyrin-G (1:1000, lab-generated^57^), goat anti-total ankyrin-G (1:1000, lab-generated ^16^), rabbit anti-KCNQ2N1 (1:500), rabbit anti-NaV (1:500, Sigma S6936), rabbit anti-neurofascin FNIII (1:500, lab generated ^58^), mouse anti-NeuN (1:1000, Sigma MAB377), mouse anti-caspr (1:1000, Neuromab 75-001), guinea pig anti-vGAT (1:1000, Synaptic Systems 131004), mouse anti-GABAA receptor β2-3 (1:1000, Sigma, MAB341), mouse anti-parvalbumin (1:1000, Sigma P3088), rabbit anti-calbindin (1:1000, Swant CB-38a). Fluorescently conjugated secondary antibodies Alexa Fluor 488, 568, or 647 (1:250, Life Technologies) and Alexa Fluor 594-Streptavidin (1:1000, Jackson ImmunoResearch 016-580-084). The following reagents were used: FBS, Poly-D-lysine, Laminin, Paraformaldehyde, DNase, Urea, and N-ethylmaleimide were from Sigma-Aldrich. B27 supplement, GlutaMAX, Penicillin-Streptomycin, Neurobasal-A, Hank’s Balanced Salt Solution, Trypsin, Hepes, Lipofectamine 2000 and Prolong Gold Antifade Reagent were from Life Technologies. Bovine serum albumin was from Gemini Bioproducts. Tween 20 was from Calbiochem.

### Image acquisition and Statistical analysis

Multi-color imaging was performed as previously described using a Zeiss 880 confocal microscope ^16^. All images were further processed in Adobe Photoshop CC 2018 software. Statistical analyses were performed using Microsoft Excel and GraphPad Prism 7. A confidence interval of 95% (P<0.05) was required for values to be considered statistically significant. All data are presented as mean ± SEM.

### Subjects, Sequencing, and Genotyping

All subjects were identified with IRB approval through the Heinz C. Prechter Bipolar Research Program at the University of Michigan. This program is the largest privately-funded longitudinal study of BD, which monitors BD patients for ten years and beyond following an initial evaluation, bi-monthly questionnaires, and neuropsychological testing by a clinical psychiatrist. The proband (II:1) was initially identified using whole genome and exome sequencing of blood samples from the Prechter Bipolar Genetics Repository at the University of Michigan. We confirmed the *ANK3* W1989R variant (rs372922084, 10:61834674 A/G) by extracting DNA from a fibroblast biopsy under IRB from the proband and performing a nested PCR initially using the primer pair 5’-GTAGCTGAAATGAAAGAGGACCT-3’ and 5’-TCTCAGAGGTGGAAGTCCTC-3’ then 5’-GATGATGAAGAACCTTTCAAAATTG-3’ and 5’-GAGGCATTTTGAGTTTGTGTTC-3’. Blood was then drawn from the mother (I:2) and father (I:1) under IRB. We extracted the DNA and performed a nested PCR using the same primers listed above. DNA from the sister (II:1) was provided from the University of Michigan Central Biorepository as part of the Heinz C. Prechter Bipolar Research Program. The 500 bp PCR fragments were purified following gel electrophoresis using a gel extraction kit and sent for sequencing at the University of Michigan DNA sequencing core. All subjects reported Caucasian ancestry.

## AUTHOR CONTRIBUTIONS

ADN, RNFC, CW, MZ, KSJ, LLS, LLI, and PMJ designed research. PMJ, KKW, KC, and VB generated and initially characterized the *Ank3* W1989R mutant mice. ADN, RNCF, JCRD, JL, KC, and CW performed experiments. ADN, RNCF, JCRD, CW, MZ, KSJ, and PMJ analyzed data. MMG provided human samples and clinical diagnosis. ADN and PMJ wrote the manuscript.

## ACKNOWLEDGEMENTS

We thank the Shanghai Radiation Facility BL19U1 and BL17U1 for X-rap beam time. Funding for this work was provided by the Michigan Predoctoral Training in Genetics (T32GM007544) (ADN), Heinz C. Prechter Bipolar Research program and Richard Tam Foundation, University of Michigan Depression Center (PMJ), R37NS076752 (LLI), National Natural Science Foundation of China (NSFC 31670734) (CW), Research Grants Council of Hong Kong (16100517 and AoE-M09-12) (MZ), and the Howard Hughes Medical Institute (VB).

